# Unifying concepts of biological function from molecules to ecosystems

**DOI:** 10.1101/105320

**Authors:** Keith D Farnsworth, Larissa Albantakis, Tancredi Caruso

**Author notes:** Corresponding author: Keith Farnsworth.

## Abstract

The concept of function arises at all levels of biological study and is often loosely and variously defined, especially within ecology. This has led to ambiguity, obscuring the common structure that unites levels of biological organisation, from molecules to ecosystems. Here we build on already successful ideas from molecular biology and complexity theory to create a precise definition of biological function which spans scales of biological organisation and can be quantified in the unifying currency of biomass, enabling comparisons of functional effectiveness (irrespective of the specific function) across the field of ecology. We give precise definitions of ecological and ecosystem function that bring clarity and precision to studies of biodiversity-ecosystem function relationships and questions of ecological redundancy. To illustrate the new concepts and their unifying power, we construct a simple community-level model with nutrient cycling and animal-plant mutualism, emphasising the importance of its network structure in determining overall functioning. This type of network structure is that of an autocatalytic set of functional relationships, which also appears at biochemical, cellular and organism levels of organisation, creating a nested hierarchy. This enables a common and unifying concept of function to apply from molecular interaction networks up to the global ecosystem.

## Function as a unifying concept for all levels of biological organisation

Are all the elements of a genome functional or are some junk? Are some species ecologically redundant or do they all have a unique role? Must a thing be naturally selected before it can be accepted as functional, or is it sufficient that it causes an effect? These questions all hinge on the definition we use for function. This became one of the key topics in a recent multi-disciplinary workshop “Functional Information: its potential for quantifying biodiversity and its relation to ecosystem functioning”, organised by the Synthesis Centre of Biodiversity Sciences, Germany. The word ‘function’ has a meaning that may initially seem self-evident and obvious, but it is used loosely and for many different meanings in biology, creating ambiguity and uncertainty which matters when we come to quantify e.g. biodiversity ecosystem function relationships. Here we draw ideas from molecular biology, community ecology, systems and information theory and philosophy of science to construct a precise and quantifiable definition of biological function that can unify and focus our thinking on the questions above. The context in which we write is one of a growing importance for functional descriptions in ecology (Krause et al., 2014; Stouffer et al., 2012), the rise of metagenomic functional analysis (Deng et al 2012; Fierer et al., 2012; Howe et al., 2014) and controversy over functional elements in, especially, the human genome (Kellis et al., 2014; Doolitle et al., 2014). Because ‘function’ is used differently among sub-disciplines of ecology and wider biology, our aim here is to promote a unifying meaning, with three goals. First, to enable cross-fertilisation of ideas through the recognition of a common concept of function, so making connections and enabling the discovery of more fundamental principles. Second, to enable communication by using ‘function’ terms in a precisely defined way. Third, to promote function as a precise, quantitative concept (with units) to establish a common framework for biodiversity-ecosystem function studies and to enable e.g. objectively based economic valuation of ecosystem components and diversity (see Farnsworth et al., 2016a). More deeply, we aim for a definition of ecological function that can illuminate the extent to which ecosystems are coherent systems, as opposed to mere arbitrary assemblies.

We start by clearly differentiating between ‘function’ and ‘trait’ in the biological context. Violle et al., (2007) dispelled much confusion over the term trait, but still today various kinds of processes and even the values of indices calculated from samples are sometimes considered ‘traits’ and this is confusing. Here, we restrict function to describe a process (an action) rather than a property of an entity. This leaves the latter to be a trait: a trait is an inherent property of a biological system (e.g. organism), which may enable a function to be performed in relation to another system (e.g. a community). For example, the specific behaviour of hiding certain plant seeds is a trait of some mammals which has the functions of dispersing the plants as well as providing a continuous food supply for the mammal. Hypothetically many more functions of this behavioural trait are conceivable, but in practice we must restrict our considerations to actual (observed) and ecologically relevant functions, as opposed to potential (hypothesised or not) functions because the full range of potential functions is indeterminate (e.g. providing food for soil invertebrates may be admissible, but providing a means of counting the mammals is not, since the latter is not an ecologically relevant function, but merely a tool of scientific observation).

The specific meaning of a function depends on both the functioning entity and the context in which it acts (function is relational, not inherent). For example a genetic element may code for a protein only in particular circumstances and a species may promote nitrogen turnover only in certain environments, therefore they only have those functions in those particular contexts. The context, in every case, is the system in which the functioning entity takes part. In a biological setting we refer to this as the next higher ‘ontological level’ recognising that living systems are composed of a nested hierarchy of organisation: systems constructed from sub-systems (Farnsworth et al., 2013). An ontological level, in this context, is a structure of biological organisation that is categorically different from those above and below in the hierarchy. For example, though a nuclear family and a whole tribe differ in organisational scale, in category both are units of human society. Conversely a collection of living cells and a human being are both different in organisational scale and different categories of entity. More formally, an ontological level (in biology) has associated with it an organisational structure with the properties of a complex system, especially the property that new phenomena emerge within it (see also Marshall et al., 2016). An ontological level is always more than a collection of objects. Thus, for example, a population (defined in the traditional sense of a mere aggregate, or count, of individuals) is not ontologically distinct from the individual organism because no properties emerge from the population level, but if the population is considered as an assembly of genes, then it may qualify as ontologically distinct from the single genome because emergent properties may appear at the level of gene-population. Further, a community may be a complex system with an organising structure that leads to emergent properties (e.g. trophic cascades or reaction-diffusion patterns), so we categorise it as a distinct ontological level. We illustrate how life as a whole can be viewed as a hierarchy of such ontological levels (Farnsworth et al., 2013) using the ontological hierarchy given in Table 2, but this does not imply that those *specific* demarkations are fixed or necessary for our understanding of function.

**Table 1.**
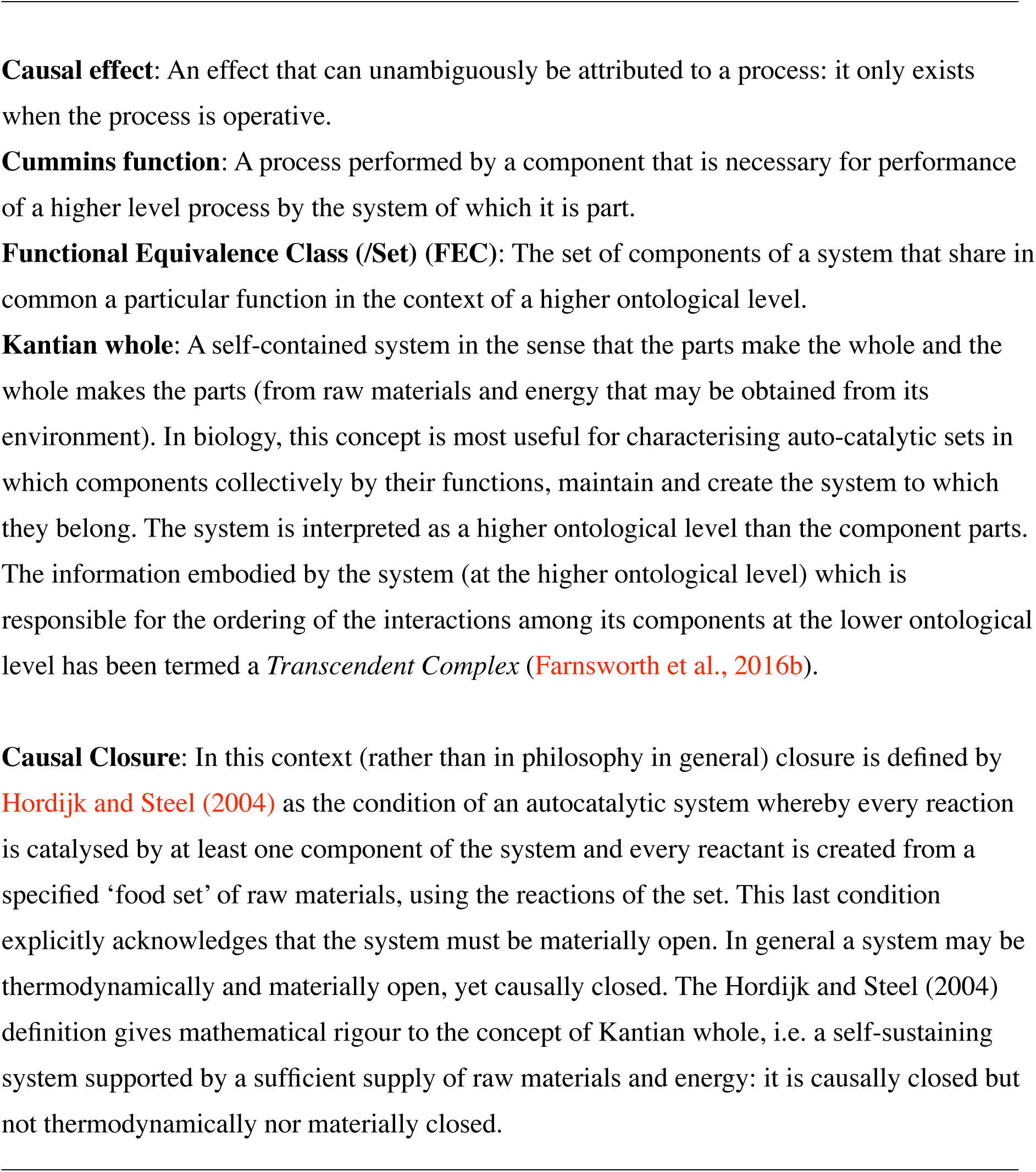
Glossary of Terms

**Table 2:**
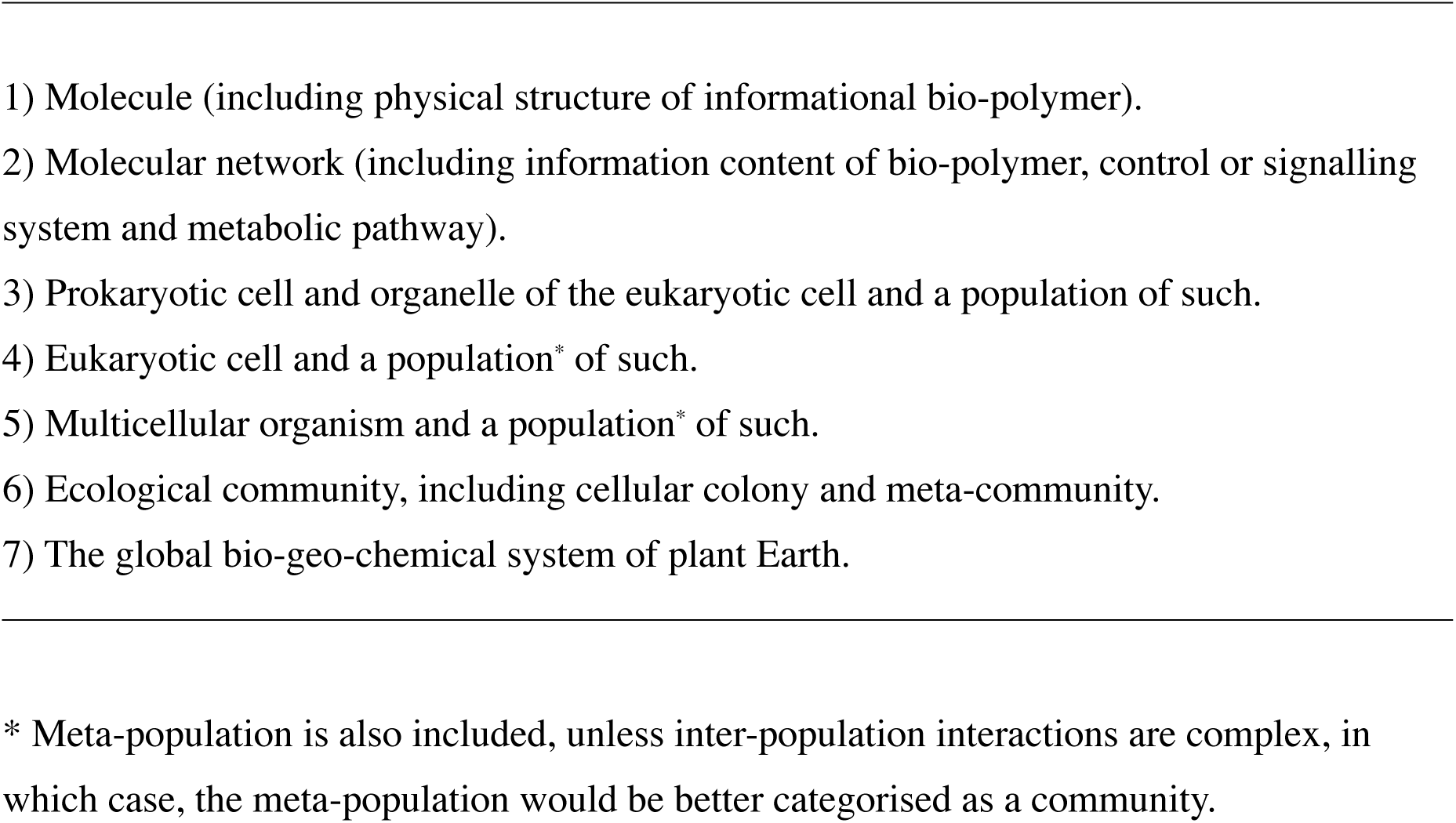
An illustrative ontological hierarchy of biological systems. At each level, properties emerge that cannot be explained by reference to its component parts (existing at the next lower level) alone. This is because every ontological level is associated with a causal interaction network among the component parts. The information embodied in this network is the source of the emergent properties.

Meta-population is also included, unless inter-population interactions are complex, in which case, the meta-population would be better categorised as a community.

From here on, we define biological function with respect to a higher ontological level. We must carefully distinguish between the terms ecological function and ecosystem function. Given our axioms, ‘ecological function’ must be an act performed by a living system *within the context of* an ecosystem. That applies equally whether the act is physiological, behavioural, one of competition, predation or mass-transfer, ecosystem engineering, or organisational - every branch of ecology can make use of the same concept here. Ecosystem function is conversely an act performed *by an ecosystem* in some ontologically higher system but what? Many ecologists seek a justification for conservation in the ecosystem services (a result of functions) provided for human society, tacitly assuming humanity to be the wider context (e.g. deGroot et al., 2012). A non-anthropocentric (objective) account places things the other way around (indeed this is a central tenet of ecological, as opposed to environmental, economics (Costanza and Daly, 1987). Objectively, the higher ontological level of ecosystem function must comprise the global geo-chemical cycle that includes the biosphere performing certain homeostatic processes (Lovelock and Margulis, 1974).

In his landmark paper, Jax (2005) resolved the use of the word ‘function’ in ecology into four broad meanings: 1) individual-level processes, such as a particular predation event; 2) systemic processes, such as nitrogen uptake; 3) individual ‘roles’ defining ‘functional groups’ of organisms, including their contribution to a higher level of organisation, such as the guild of detritivores, or simply a phenotypic (often life-history) category and 4) effects of the activity of working ecosystems that impinge on human society, leading to ecosystem services. We see immediately that (3) describes a set of functional capabilities of organisms and (1) describes the realisation of one or more of these in practice, whilst (2) does the same but takes a coarse-grained view of the system and (4) interprets this from an anthropocentric perspective.

In the following, using examples from ecology and molecular biology, we will argue for a more constrained definition of function, in line with (3): that a biological function must demonstrate causal effect (Table 1) from one to a higher ontological level of biological organisation. This relationship enables us to identify ‘functional equivalence sets’ (Table 1), shifting the focus from particular systems, defined by their traits, to functional classes. This move enables a logically self-consistent and integrative way to assess all kinds of interaction in biological systems, by starting from a network of functional equivalence sets. Only by doing this can we be sure that function has a standard and inter-comparable meaning, in for example biodiversity-function relationships.

## Insights from philosophy and molecular biology

A system composed of parts, each of whose existence depends on that of the whole system is termed a ‘Kantian whole’ (Table 1), the archetypal example being a bacterial cell (Kauffman and Clayton 2006). The concept of function arises to describe the role that the parts play in the system, without having to descend into non-scientific teleology. (Note - the origin of this terminology lies in Emanuel Kant’s definition of an organised whole (Ginsborg, 2006)). Accordingly, the philosopher Cummins (1975) proposed that ‘function’ is an objective account of the contribution made by a system’s component to the ‘capacity’ of the whole system. Thus *Cummins function* (Table 1) describes a relationship between a system and one or more of its component parts. The relationship is that at least one process performed by the component/s is *necessary* for a process performed by the whole system, as in the case of category (3) of the Jax (2005) schema. This is a powerful idea we shall use from here on. What we add is clarification about what specifies the ‘whole system’, particularly how it is bounded, and how this concept can be turned into a tool for quantifying function, especially, expanding the notion of being ‘necessary’ to a rigorous quantification of causal effect. Natural selection for a particular component within an organism provides a clear example. In that case, the component (e.g. an eye) contributes a process that increases the (quantifiable) Darwinian fitness of the organism, thereby demonstrating Cummins function. The idea, that ‘true biological functions’ should be naturally selected, has already emerged in the field of molecular biology, though a broader set of function is advocated by some - as we next discuss.

Recently, thousands of full genomes have been sequenced, not the least being the human genome, and efforts to interpret this new data have stirred up the controversy over ‘junk DNA’, which is taken to be the alternative to ‘functional DNA’ (see Kellis et al., 2014). In this field, little attention has been paid to the precise meaning of ‘function’, despite much loose talk of ‘functional DNA’. In their criticism, Doolitle et al., (2014) clarified the range of potential meanings of the term and the conceptual errors that may arise from failing to resolve them. The most justifiable assignment of function to a DNA element, they argued, was where selection at the organism level was demonstrated. This is only a subset of cases where a causal role has been established. Alternatives include a) selection at higher or lower ontological levels, b) neutral (non-selective) processes, which frequently ratchet their way into conserved stability, and c) spandrels (the term used by Gould and Lewontin (1979), for by-products of selection). Doolitle et al. (2014) were careful to distinguish the spandrels from “mere effects”, which play no causal role in the system to which they belong: spandrels play an unintended and perhaps irrelevant causal role (e.g. the ‘thumping sound’ of the heart).

Building on this, we may at least ascribe the word ‘functional’ to biological systems having causal effects that are known to have been naturally selected for at any level of organisation subject to natural selection (not only species, but also gene, or genetic network). They are functional in the Cummins sense and their causal effect can be quantified by the change in fitness they cause in the system to which they belong. However, this definition would leave out systems with Cummins function but not derived from natural selection. They would be left as ‘merely’ causal, rather than biologically functional. These causal systems could be subsystems of naturally selected systems, for example ‘molecular machines’ (e.g. the ATP synthase complex), or biochemical networks (examples in Jaeger and Calkins, 2012), or they may be super-systems of them: populations or communities of organisms.

Metagenomic functional analysis challenges this restrictive notion of biological function by revealing the processes that are being (RNA), or could be (DNA) performed at the community level of organisation (Tringe et al., 2005; Warnecke et al., 2007; Huttenhower et al., 2012; Howe et al., 2014). With it, function is readily discerned because the link between gene expression and functional protein or peptide is usually known and unambiguous, irrespective of our knowledge of its natural selection. Although eukaryotic organisms typically present more complicated and often less clear geno-phenotypic links, the method is also developing for them (see e.g. Knack et al., 2015). These new molecular tools reveal a literal functioning: the causal chain leading from one biomolecule to the next. The components (e.g. functional genes) are performing Cummins function for the organism level (which because of this, performs Cummins function at the community level) and their performance is well defined and quantifiable. This unambiguous identification of a causal chain is the ideal towards which ecologists may strive (Gotelli et al., 2012; Bohmann et al., 2014). Accordingly, we now offer a definition for biological function:

## A biological function is a process enacted by a biological system A at ontological level *n* which influences one or more processes of a system B at level *n*+1, of which A is a component part

This definition is similar to that of Cummins function, but more precisely identifies the link between ontological levels of organisation. By tracing functional effects through the nested hierarchy leading from one to the next higher ontological level, it can be made explicit how a function at the molecular level can be functional at an ecological level. The definition does not require a history of natural selection and it does not require a ‘good’ outcome for the system (as selection would). The latter point answers the philosophical objection often levelled at scientific accounts of function (see e.g. Griffiths, 1993), which arises from the teleological (referring to purpose) and normative (what ought to be) connotations of the every-day use of the word ‘function’ (a rich literature on philosophical approaches to biological function is reviewed by Neander (2011)). Our definition creates a clear separation between these everyday meanings of function and a strict scientific meaning.

If functions are strictly processes, then potentially more than one system component can perform them and functional redundancy and substitution become possible. This idea, which we now incorporate into our definition of function is usefully formalised by the concept of *functional equivalence class* (FEC) (Table 1), which has grown from analysis of biochemical networks. The FEC was defined by Auletta et al., (2008) as a set of biochemical ‘operations’ having effects in common which are relevant to ‘goals’. The FEC consists of all operations (behaviours or processes) having the effect in question and this is context-dependent because an effect always depends on the nature of both the subject and the object. For example, the DNA sequences and corresponding protein structures of alcohol dehydrogenases in vertebrates bear no similarity with those of Drosophila and they work through different chemical reactions, but achieve the same end result of removing hydrogen from alcohol (Doolitle 1994). Those different dehydrogenasing processes form an FEC with respect to frugivorous organisms (and would not with respect to obligate carnivores). The existence of alcohols in fruit arises from interactions among plants and micro-organisms (i.e. at the community level) and exerts natural selection on the metabolic processes (cellular level) of frugiverous organisms. This illustrates a general feature of FECs: they define the function at one ontological level in terms of processes at a higher level, which in turn must include at least one component from among them. Any function specifying an FEC complies with the definition of Cummins function. Using the FEC generalises the concept by identifying the source of causation: it is not the individual components performing the function, but the FEC (i.e. the macro-level unit). This allows for the possibility of functional redundancy and substitution: both important features illustrated at the ecological level by species turn-over and at the sub-species level by heterologies and convergence (reviewed by McGhee, 2011). We next discuss the extent to which this definition of biological function applies in ecology, with special reference to the biodiversity-function relationship.

### At what level do whole ecosystems function?

Function in ecological communities had traditionally been thought different from function in whole or parts of organisms because ecological systems do not fit the conventional model of Darwinian evolution (Maclaurin and Sterelny, 2008). Recent evidence (e.g. Rillig, et al., 2015) and an emerging ecological theory of broader evolution (Laland et al., 2015)challenges that view. Niche construction theory (Odling-Smee et al. 1996) and the concept of reciprocal causation in which organisms influence their evolution via ecological processes (Laland et al., 2011) both include the ecosystem as an integral part of a complex evolutionary process.

The predominance in community ecology of predator-prey networks, described by flows of energy in foodwebs has emphasised an incomplete model of community structure in which only one function (energy flow) is admitted, rendering what Loreau (2010) calls ‘horizontal diversity’ redundant. In every real community, each organism performs more than one function and functions are more than contributions to energy flow. This has been recognised in the development of community models based on nutrient-cycling (e.g. Loreau, 1996; Thébault and Loreau (2003); Loreau (2010) and in the study of mutualistic networks involving pollinators and seed dispersal (e.g. Schleuning et al., 2015), to which parasites and ‘ecosystem engineers’ might be added (see e.g. Bruno et al., 2003). As Krause et al., (2014) explicitly state, ecosystem functioning results from the interactions among organisms, which though based on organism diversity are a (neglected) form of diversity in themselves. Our definition of function as strictly relational calls attention to the importance of the interaction network as a component of diversity and it identifies the organisation (network structure) of its component ecological functions as the (proximate) cause of ecosystem function. This is a point we wish to emphasise, especially because so far, the mainstream of studies into biodiversity / ecosystem function (BEF) relationships has neglected the contribution of the community network (in and of itself) to ecological functioning (Hooper et al., 2005; Cardinale et al., 2012), despite contradictory evidence (e.g. Bascompte, 2009; Grey et al., 2014; Valiente-Banuet et al., 2015). Experimental manipulations that discriminate the effect of network structure (the organisation of functions at the community level) from organism effects (the traits enabling ecological functions) will be challenging, but are necessary to appreciate the extent to which the higher ontological level of community operates.

### What this says about functional redundancy

Functional redundancy does not mean that a system will be unaffected by species loss. The idea that species may be mutually redundant can arise from the neglect of the quantitative contributions made by *individual* organisms to ecological processes. Functionally substitutability (belonging to the same FEC) does not imply mutual redundancy: an analogy with members of a tug-of-war team illustrates how individual organisms can be functionally substitutable, but not redundant. In community ecology, this point is well illustrated by empirical findings, e.g. from O’Gorman and Emmerson (2009) and Isbell et al., (2011). Further, we must distinguish between redundancy and degeneracy. Consider a set of n species <u>S</u>, and a set of k functions <u>F</u>. We associate a *k*-long vector of functions <u>f</u>_i_ with each member species s_i_, the values in the vector being the quantified contributions of s_i_ to each of the k functions in <u>F</u>. Now we can transform from n species to m functional equivalence sets <u>E</u> by gathering all those members of <u>S</u> sharing the same <u>f</u>. But being relational, functional equivalence must be defined relative to the context, i.e. the environmental conditions. Organisms that are members of FEC e_j_ in <u>E</u>, irrespective of environment are qualitatively redundant, but those that only share <u>f</u>_j_ in common for a particular environment are merely degenerate with respect to that environment alone (Tononi et. al., 1999). This is the basis for the ‘insurance’ justification for biodiversity in the environmental economics literature (Yachi and Loreau, 1999; Baumgärtner, 2007).

Community ecology and especially BEF research has tended to account for one single species-level Cummins function at a time (e.g. total energy flow), though this constriction was eased, notably by Gamfeldt et al., (2008) and Isbell et al., (2011) and by attempts to integrate mutualistic networks along with competition, predator prey and parasitism (see Kefi et al., 2012 and reviews by Krause et al., 2014 and Namba, 2015). Since each organism can perform many functions and each function can be performed by many organisms, it makes sense to transform from networks of species-specific populations to quantitative functional networks of FECs. Many community ecologists have started to refocus this way (e.g. Schleuning et al., 2015), aided by increased data from molecular ecology which can more precisely isolate function (e.g. Zhou et al., 2010; Deng et al., 2012; Fierer et al., 2012). There is insufficient evidence to claim that any organism is functionally redundant in the quantitative, multi-functional sense (Gamfeldt et al., 2008, Farnsworth et al., 2012) and we think it unlikely.

## Quantifying function

Identifying FECs is a key component of the functional analysis of biological systems. FECs may be equated with observed functions from metagenomic functional analysis (see e.g. Fierer et al., 2012) and other identified processes such as metabolism, predation and specific ecosystem engineering. In practice, this may result in too many classes for quantitative analysis. We would then be forced (as we usually are) to identify a set of *a-priori* function classes to which observed functions are assigned. Choosing these classes will depend on what degree of difference among functions we would consider sufficient to identify them as separate. This may be objectively estimated from the overlapping of the functions’ effects, for which the concept of ‘effective number’ in biodiversity (Jost, 2006, Chao et al., 2014) was applied to ecosystem processes by Ulanowicz et al., (2014). Observed functions would be assigned in proportion to both the number of components (enzymes, populations, etc.) and the effectiveness of the functions in any given environment. Effectiveness is empirically quantifiable given a precise definition of the function as a process. For some processes this approach is already well established. For example, predation is quantified by the individual (per-capita) ‘functional response’ (see e.g. Dick et al., (2014) for a relevant application). Ecosystem engineering is often highly context specific, but empirical approaches have been found (e.g. Queiros et al., 2011). The approach we are suggesting calls for a precise definition and identification of FECs, for which the payoff is likely to be a more robust understanding of how biological systems work.

As we have defined them, functions are causal. Thus in principle we can quantify them in both magnitude and direction, according to their causal effect (for which structural equation modelling and related methods may be useful (Shipley, 2000). In practice this can be very difficult for complex systems with reciprocal causation (Laland et al., 2011), particularly in the face of limited time-series data samples, latent variables, and incomplete knowledge of the system under consideration ( see Sugihara et al., 2012 and Deyle et al., 2016). Recent developments in information-theoretic and network analysis (e.g. Zenil et al., 2016) combined with knock-out experiments to discern causal interactions (Pearl, 2000) provide a way forward. Measures of functional effectiveness can, in principle, be assessed at and across different levels of organisation of a system (Marshall et al., 2016), but application of these rigorous theoretical measures to ecological systems demands the acquisition of extensive time-series data.

As a specific example consider the (highly simplified) illustrative model community in Figure 1 (detailed description in Appendix), which depicts four trophic levels and five functions: nitrogen supply (recycling, fixation, nitrification etc.), carbon supply, with the consequent functions of carbon and nitrogen flow, and reproductive facilitation (e.g. pollination). Note that the functions are individually quantified in their own *native* units (i.e. units of carbon flow rate, nitrogen flow rate and e.g. pollination rate). Every function performed in the system is contingent on every other, because each depends on the biomass and thereby the growth rate of the species in each FEC providing it. This in turn depends on the species supplying functions necessary for this growth. Analysis of functional dependencies (Figure 2) shows that every function ultimately depends on the aggregate community-level biomass production rate (B), but this in turn depends on every function. This illustrates two important principles: first that the FECs form an autocatalytic set (*sensu*
Kauffman, 1986) and second that due to the recursive structure (everything depends on everything else) the system is in causal closure (Table 1) and thereby (Kauffman and Clayton, 2006) constitutes a Kantian whole. The closure referred to here is strictly causal and not material, nor thermodynamic. Causal closure means that the network of causal interactions is sufficient to describe the behaviour at the system level, given the necessary material and energy supply from the environment (Hordijk & Steel, 2004), acknowledging that living systems are always materially and thermodynamically open. This causal closure is a general property of the physiology of individual organisms (Luisi 2003), many biochemical sub-systems of life (Kauffman 1986) and as our example illustrates, at least some ecological systems (for which it has important implications). As an illustration of the mutual dependencies, Figure 3 shows the equilibrium biomass (excluding nutrient providers) of the community which varies with a) the asymptote of nutrient providers’ functional response and b) the effectiveness of reproduction facilitation for plants by animals. The community dynamics are tremendously complicated (Appendix) as is typical of a system with several autocatalytic loops and the result of exogenous perturbations is determined by internal relationships and correspondingly difficult to predict.

**Fig 1.**
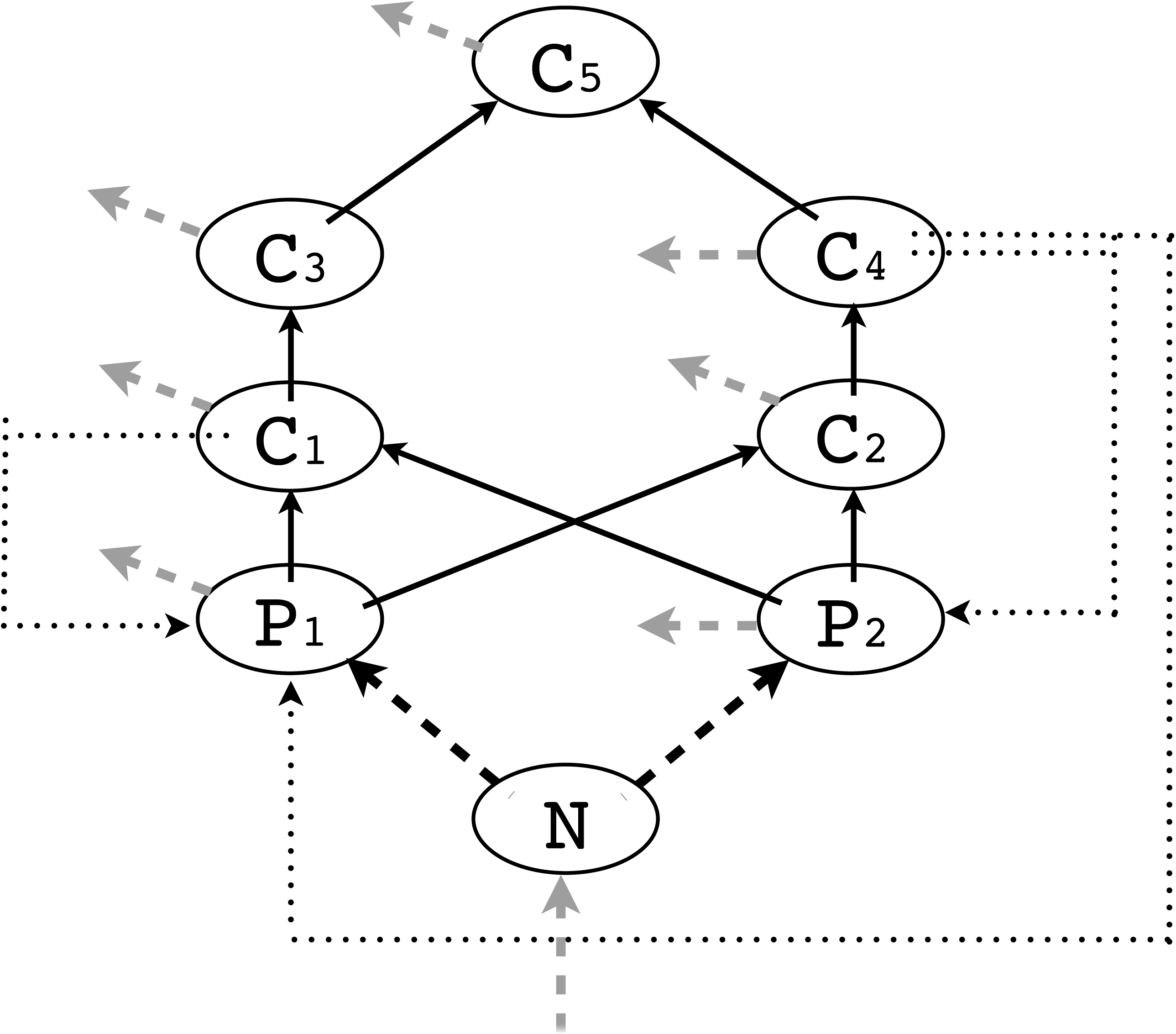
An illustrative ecological community consisting of primary producer FECs (P_1_, P_2_); primary consumers (C_1_,C_2_), secondary consumers (C_3_,C_4_ and C_5_) and nitrogen suppliers (N). Thin solid arrows show trophic flows, thin dotted arrows show reproductive facilitation (e.g. pollination and seed dispersal), thick black dashed arrows show nutrient flow (in isolation) and grey dashed show waste nitrogen, which is assumed all to return to N. The system can be envisaged as three parallel functional networks (nutrient recycling, carbon pumping and reproductive facilitation). Together, these functions catalyse one another so that the whole system is an autocatalytic-set.

**Fig. 2.**
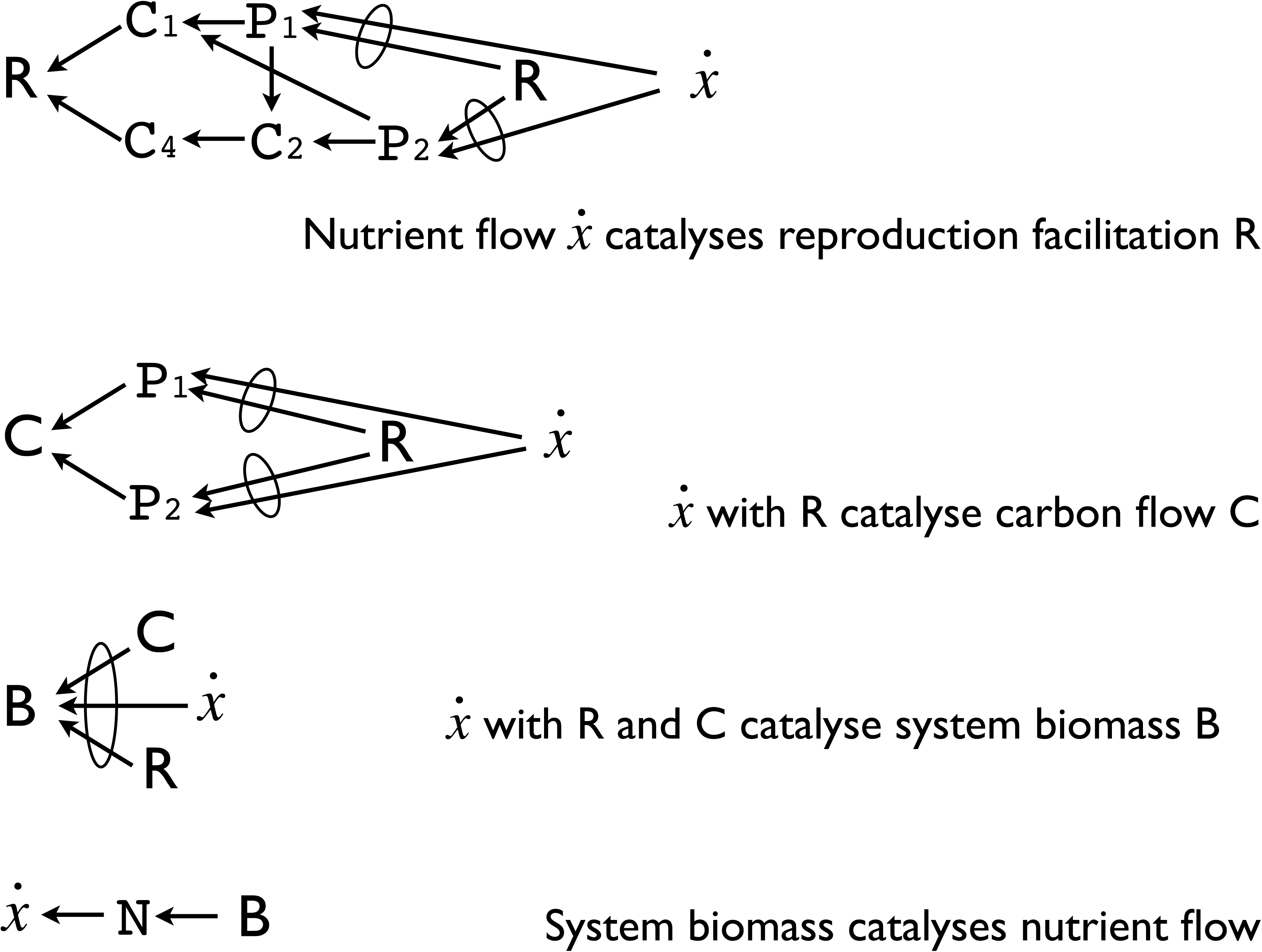
Functional dependencies show the community to be an autocatalytic set. In (a) reproductive facilitation (R) depends on FECs C_1_ and C_4_, which in turn depend on biomass production of P_1_ and C_2_, respectively, which in turn depend on nitrogen supply from N and reproductive facilitation R - forming an autocatalytic loop. The ellipse symbols indicate ‘jointly necessary’, otherwise parallel inputs are mutually degenerate, or redundant. C: carbon supply; N: nitrogen supply, B(E) is biomass production rate of the whole community. Note that all functions ultimately depend on - and result in - B(E). For this reason, B(E) provides a suitable ‘master function’ with which to quantify all internal functions.

**Fig. 3.**
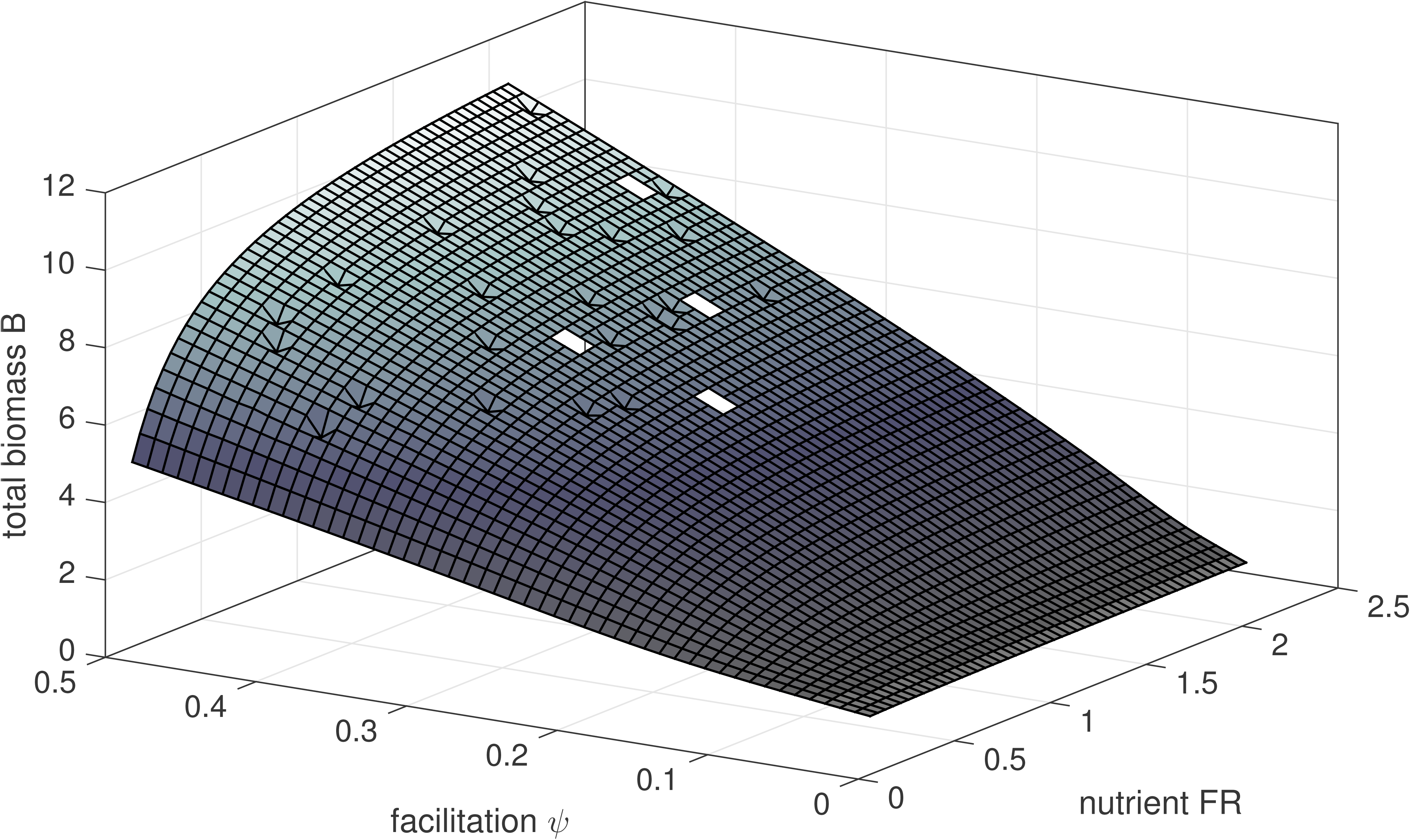
How equilibrium total system biomass B - the master function - varies with the rate of plant reproductive facilitation by consumer species and also the asymptote of the functional response of nutrient providers, to supply of raw materials (nutrient FR). The plot was formed from finite-difference numerical calculations of the system (Appendix equations 6-13). The dynamics did not converge to an equilibrium for combinations of values, which is the reason for the few missing and aberrant results (intentionally left visible in the plot).

### The concept of Master Function

Most broadly, a community that can be viewed as a Kantian whole is consistent with a general definition of life: “*A system can be said to be living if it is able to transform external matter and energy into an internal process of self-maintenance and production of its own components*” (Luisi, 2003, p52). By this definition, all biological function ultimately amounts to production of living cells and this is readily quantified by biomass. Accordingly, we can regard biomass production as a ‘master function’ (Jaeger and Calkins, 2012), meaning that all functions may be quantified by their contribution to it (by analogy, all processes in a factory may ultimately be quantified by their contribution to the profit made by the factory owning business). Alternatives to biomass have been inspired by approaches from physics and computer science: energy dissipation based on thermodynamic theory, e.g. Kaila and Annila (2008) and diversification based on complexity theory (Kauffman, 2000). We chose to focus on biomass production because it is most readily measured in practice and also has a clear biological and evolutionary basis. The idea of a master function removes the need for a teleological account of function: it is the de-facto end point of the causal chains to which all other biological functions belong, in the proximal as opposed to ultimate sense of Mayr’s (1961) dichotomy (modified by Laland et al., 2011). Any biological process which enhances the fitness of a biological system, increases the system’s potential to reproduce and by this, over time, it increases the amount of biomass embodying it. Using biomass as the ‘master function’ means that all functions at all scales of biological organisation can be quantified in terms of their contribution to biomass production (which may be found *in vivo* by e.g. species removal, or gene knockouts, or in computational simulations through sensitivity analysis as illustrated in Figure 3). So whilst every kind of function can be quantified in its own native units, we can also quantify it in the community context (referring to the function to which it contributes at the higher ontological level), by its effect on aggregated biomass production. Notice that this is true at the community level, but also at the level of cells, where reproduction via completing the cell-cycle is the ‘master-function’ (as illustrated in Jaeger and Calkins, 2012). In our suggested interpretation, biomass production is a universal currency for biological function, which integrates ecological function with that defined at any other level of biological organisation. We can quantify function (of any process) in terms of the specific change in the rate of creation of cells (fitness), given the presence / absence of the functional component (this is analogous to the power density proposed by Chaisson (2011) as a surrogate for complexity). While individual processes of course all appear in different units, the integrative framework proposed enables them all to be interpreted in the common currency of biomass production, using e.g. mathematical simulations to examine the effect of removing any functional component upon the production rate of the remaining system, following recovery to a new equilibrium (as in Fung et al., 2015).

## Conclusion

The Biblical lesson of the ‘tower of Babel’ was that division and confusion results from a fragmentation of language. By establishing a common and precise meaning for the word ‘function’, we believe a conceptual unity can arise among the various sub-disciplines of biology. We have found that biological systems at all levels of organisation share the properties of auto-catalytic sets and that this enables us to summarise the aggregate effect of ecological functions via specific biomass production rate, which serves as an integrating surrogate measure of their effectiveness. Specifically, to quantify biological function, irrespective of whether this is genetic, cellular or ecological (behavioural, physiological, population, community or process), we suggest the following steps:

1) Identify the ontological level of the system under study and the next higher ontological level of which it is a component (Table 2).
2) Identify the FEC(s) to which the study system belongs (in relation to the higher ontological level) - this amounts to specifying the system’s functions in terms of relationships among the parts.
3) Perform knockout / elimination experiment (or simulation study) to quantify function in terms of the master function for the higher ontological level (the whole of this level should be included in the experiment).

This approach is now well established in molecular biology for investigating gene regulation networks, the result of protein expression and signalling networks. It is developing in the analysis of prokaryotic communities via functional analysis of ‘environmental’ gene sequence data. Here we have shown that it can be extended to levels of organisation beyond the individual organism and applied generally in all fields of ecological science, thus integrating the concept of function and its analysis over all levels of biological organisation.

## Acknowledgements

This work was derived from discussions held at the Synthesis Centre for Biodiversity Sciences (sDiv), which inspired and funded the meeting. L. Albantakis received support from the Templeton World Charities Foundation Grant TWCF 0067/AB41.

## References

Auletta, G., et al. 2008. Top-down causation by information control: From a philosophical problem to a scientific research programme. - J. R. Soc. Interface 5, 1159–1172.

Bascompte, J. 2009. Disentangling the Web of Life. - Science 325: 416–419.

Baumgärtner, S. 2007. The insurance value of biodiversity in the provision of ecosystem services. - Nat. Resour. Model. 20:87–127471.

Bohmann, K.A. et al. 2014. Environmental DNA for wildlife biology and biodiversity monitoring. - Trends Ecol. Evol. 29:358–367.

Bruno, J. F. et al. 2003. Inclusion of facilitation into ecological theory. - Trends Ecol. Evol 18, 119–125.

Cardinale, B. J., et al. 2012. Biodiversity loss and its impact on humanity. - Nature 486, 59–67.

Chaisson, E. J. 2011. Energy rate density as a complexity metric and evolutionary driver. - Complexity 16:27–40.

Chao, A., et al. 2014. Unifying Species Diversity, Phylogenetic Diversity, Functional Diversity, and Related Similarity and Differentiation Measures Through Hill Numbers. - Ann. Rev. Ecol. Evol. S. 45:297–324.

Costanza, R. and Daly, H. E. 1987. Toward an ecological economics. - Ecol. Model. 38:1–7.

Cummins, R. 1975. Functional analysis. - J. Philos. 72(20):741–765.

De Groot, R.S., et al. 2012. Global estimates of the value of ecosystems and their services in monetary units. - Ecosyst. Ser v. 1, 50–61.

Deng, Y., et al. 2012. Molecular ecological network analyses. - BMC Bioinformatics 13:113.

Deyle, E. R. et al. 2016. Tracking and forecasting ecosystem interactions in real time. - Proc R., Soc. B-Biol. Sci. 283(1822).

Dick, J. T. A. et al. 2014. Advancing impact prediction and hypothesis testing in invasion ecology using a comparative functional response approach. - Biol. Invasions 16:735–753.

Doolitle, R. F. 1994. Convergent evolution: the need to be explicit. - Trends Biochem. Sci 19:15–18.

DoolittleW. F. et al. 2014. Distinguishing between "Function" and "Effect" in genome biology. - Genome Biol. Evol. 6: 1234–1237.

Farnsworth, K. D. et al. 2012. Functional Complexity: The source of value in biodiversity. - Ecol. Complex. 11: 46–52.

Farnsworth, K. D. et al. 2013. Living is Information Processing: From Molecules to Global Systems. - Acta Biotheor. 62: 203–222.

Farnsworth, K.D. et al. 2016a. The complexity of biodiversity: A biological perspective on economic valuation. - Ecol. Econ. 120, 350–354.

Farnsworth, K. D. et al. 2016b. Living through downward causation: from molecules to ecosystems, in S.I. Walker, G.F.R. Ellis and P.C.W. Davies (eds) From Matter to Life: Information and Causality. - Cambridge University Press.

Fierer, N. et al. 2012. Cross-biome metagenomic analyses of soil microbial communities and their functional attributes. - Proc. Natl. Acad. Sci. U. S. A. 109, 21390–21395

Fung, T. et al. 2015. Impact of biodiversity loss on production in complex marine food webs mitigated by prey-release. - Nat. Comm. 6. (no page numbers with this journal)

Gamfeldt, L. et al. 2008. Multiple functions increase the importance of biodiversity for overall ecosystem functioning. - Ecology 89:1223–1231.

Ginsborg, H. 2006 Kant’s biological teleology and its philosophical significance. In Bird, G(Ed.) A Companion to Kant. - Wiley-Blackwell

GouldS. J.and LewontinR. C. 1979. The spandrels of San Marco and the Panglossian paradigm: a critique of the adaptationist programme. - Proc R. Soc. B. 205:581–598.

Gotelli, N. J. et al. 2012. Environmental proteomics, biodiversity statistics and food-web structure. - Trends Ecol. Evol. 27:436–442.

Grey et al. 2014. Ecological networks: the missing links in biomonitoring science. - J. App.Ecol. 51:1444–1449.

Griffiths, P. E. 1993. Functional analysis and proper functions. - Brit. J. Philos. Sci., 44:409–422.

Hooper, D.U. et al. 2005. Effects of biodiversity on ecosystem functioning: a consensus of current knowledge. - Ecol. Monogr. 75, 3–35.

Hordijk, W. and Steel, M. 2004. Detecting autocatalytic, self-sustaining sets in chemical reaction systems. - J. Theor. Biol. 227, 451–461

Howe, A. C. et al. 2014. Tackling soil diversity with the assembly of large, complex metagenomes. - Proc. Natl. Acad. Sci. U. S. A. 111:4904–4909.

Huttenhower. et al. 2012. Structure, function and diversity of the healthy human microbiome. - Nature 486:207–214.

Isbell, F. et al. 2011. High plant diversity is needed to maintain ecosystem services. - Nature 477:199–202.

Jax, K. 2005. Function and “functioning” in ecology: what does it mean? - Oikos 3:641–648.

Jaeger, L. and Calkins, E. 2012. Downward causation by information control in micro-organisms. - Interface Focus 2, 26–41.

Jost, L. 2006. Entropy and diversity. - Oikos 113:363–75.

Kaila, V.R.I. & Annila, A. 2008. Natural selection for least action. - Proc. R. Soc. A 464, 3055–3070.

Kauffman, S. A. 1986. Autocatalytic sets of proteins. - J. Theor. Biol. 119:1–24.

Kauffman, S. A. 2000. Investigations. - Oxford University Press.

Kauffman, S. A. and Clayton, P. 2006. On emergence, agency and organisation. - Phil. Biol 21:501–521.

Kefi, S.. et al 2012. More than a meal … integrating non-feeding interactions into food webs. - Ecol. Lett. 15:291–300.

KellisM. et al. 2014. Defining functional DNA elements in the human genome. - Proc. Natl Acad. Sci. U S A. 111:6131–6138.

Knack, J. J.. et al. 2015. Microbiomes of streptophyte algae and bryophytes suggest that a functional suite of microbiota fostered plant colonization of land. - Internat. J. Plant Sci. 176:405–420.

Krause, S. et al. 2014. Trait-based approaches for understanding microbial biodiversity and ecosystem functioning. - Front. Microbiol 5. Article 251

Laland, K.N.. et al. 2015. The extended evolutionary synthesis: its structure, assumptions and predictions. - Proc. R. Soc. B. 282: 2015101

Laland, K. N.. et al. 2011. Cause and Effect in Biology Revisited: Is Mayr’s Proximate-Ultimate Dichotomy Still Useful? - Science 334.6062 : 1512–1516.

LoreauM. 1996. Coexistence of multiple food chains in a heterogeneous environment: interactions among community structure, ecosystem functioning, and nutrient dynamics. - Math. Biosci. 134: 153–188.

Loreau, M. 2010. From populations to ecosystems: Theoretical foundations for a new ecological synthesis. - Princeton University Press.

Lovelock, J. E. and Margulis, L. 1974. Atmospheric homeostasis by and for the biosphere:The Gaia hypothesis. - Tellus 26:2–10.

Luisi, P. L. 2003. Autopoiesis: a review and a reappraisal. - Naturwissenschaften 90:49–59.

Marshall, W.. et al 2016. Black-boxing and cause-effect power. - arXiv 1608. 03461.

Maclaurin, J. and Sterelny, K. 2008. What Is Biodiversity? - Univ. Chicago Press.

Mayr, E. 1961. Cause and effect in biology. - Science 134: 1501.

McGhee, G. R. 2011. Convergent evolution: limited forms most beautiful. - M.I.T Press

Namba, T. 2015. Multi-faceted approaches toward unravelling complex ecological networks. - Popul. Ecol. 57:3–19.

Neander, K. 2011. Routledge Encyclopedia of Philosophy (Online). - Routledge

O’Gorman, E. J. and Emmerson, M. C. 2009. Perturbations to trophic interactions and the stability of complex food webs. - Proc. Natl. Acad. Sci. U. S. A. 106 (32), 13393–13398.

Odling-Smee, F.J.. et al. 1996. Niche construction. - Am. Nat. 147, 641–648.

Queiros, A. et al. 2011. Context dependence of marine ecosystem engineer invasion impacts on benthic ecosystem functioning. - Biological Invasions. 13:1059–1075.

PearlJ. 2000. Causality: models, reasoning and inference. - Cambridge Univ. Press.

Rillig, M.C. et al. 2015. Interchange of entire communities: microbial community coalescence. - Trends Ecol. Evol., 30, 470–476.

Schleuning, M. et al. 2015. Predicting ecosystem functions from biodiversity and mutualistic networks: an extension of trait-based concepts to plant-animal interactions. - Ecography 38:380–392.

Shipley, B. 2000. Cause and Correlation in Biology. - Cambridge Univ. Press.

Stouffer, D. B. et al. 2012. Evolutionary Conservation of Species’ Roles in Food Webs. - Science. 335:1489–1492.

Sugihara, G. et al. 2012. Detecting Causality in Complex Ecosystems. - Science 338:496–500.

ThébaultE., Loreau, M. 2003. Food-web constraints on biodiversity-ecosystem functioning relationships. - Proc. Natl. Acad. Sci. USA 100:14949–14954.

Tringe, S. G. et al. 2005. Comparative metagenomics of microbial communities. - Science 308:554–557.

TononiG. et al. 1999. Measures of degeneracy and redundancy in biological networks. - Proc. Natl. Acad. Sci. U. S. A. 96: 3257–3262.

UlanowiczR. E. et al. 2014. Limits on ecosystem trophic complexity: insights from ecological network analysis. - Ecol. Lett. 17: 127–136.

Valiente-Banuet, A.. et al. 2015. Beyond species loss: the extinction of ecological interactions in a changing world. - Funct. Ecol. 29:299–307.

Violle, C.. et al. 2007. Let the concept of trait be functional! - Oikos. 116:882–92.

Warnecke, F. et al. 2007. Metagenomic and functional analysis of hindgut microbiota of a wood-feeding higher termite. - Nature. 450:560–U17.

Yachi, S. and M., Loreau. 1999. Biodiversity and ecosystem productivity in a fluctuating environment: The insurance hypothesis. - Proc. Natl. Acad. Sci. U. S. A. 96:1463–1468.

Zenil, H.. et al. 2016. Methods of information theory and algorithmic complexity for network biology. - Seminars in Cell & Developmental Biology 51: 32–43.

Zhou, J. Z. et al. 2010. Functional Molecular Ecological Networks. - mBio 1. (there are no pages numbers with this online journal)

